# SPAs promote thermomorphogenesis via regulating the phyB-PIF4 module in *Arabidopsis*

**DOI:** 10.1101/2020.02.07.938951

**Authors:** Sanghwa Lee, Inyup Paik, Enamul Huq

**Affiliations:** Department of Molecular Biosciences and The Institute for Cellular and Molecular Biology, The University of Texas at Austin, Austin, Texas 78712

**Keywords:** Arabidopsis, high ambient temperature, SUPPRESSOR OF PHYA-105 (SPA), PIF4, phytochrome B, thermomorphogenesis

## Abstract

High ambient temperature due to global warming has a profound influence on plant growth and development at all stages of life cycle. Plant response to high ambient temperature termed thermomorphogenesis is characterized by hypocotyl and petiole elongation, and hyponastic growth at seedling stage. However, the molecular mechanism of thermomorphogenesis is still rudimentary. Here, we show that a set of four *SUPPRESSOR OF PHYA-105* (*SPA*) genes are required for thermomorphogenesis. Consistently, SPAs are necessary for global gene expression changes in response to high ambient temperature. SPA1 level is unaffected, while the thermosensor phyB is stabilized in the *spaQ* mutant at high ambient temperature. Furthermore, in the absence of four *SPA* genes, the pivotal transcription factor PIF4 fails to accumulate, indicating a role of SPAs in regulating the phyB-PIF4 module at high ambient temperature. SPA1 directly phosphorylates PIF4 *in vitro*, and a mutant SPA1 affecting the kinase activity fails to rescue the PIF4 level as well as the thermo-insensitive phenotype of *spaQ*, suggesting that the SPA1 kinase activity is necessary for thermomorphogenesis. Taken together, these data suggest that SPAs integrate light and temperature signaling via fine tuning the phyB-PIF4 module.

## INTRODUCTION

Among many facets of climate change, global warming causing an elevated ambient temperature has a profound influence on plant growth and development at all stages of life cycle (Lippmann et al., 2019). Plants modify their body plan as an adaptive response to cope with the high ambient temperature. These suite of growth and developmental changes are termed thermomorphogenesis, which is characterized by hypocotyl and petiole elongation, hyponastic growth, reduced stomatal density and early flowering (Casal and Balasubramanian, 2019; Quint et al., 2016).

Recent reports showed that the temperature and light perception mechanisms are tightly linked. For example, the red light photoreceptor phytochrome B (phyB) and the blue light photoreceptor Cryptochrome 1 (Cry1) have been shown to function as thermosensors (Jung et al., 2016; Legris et al., 2016; Ma et al., 2016). Downstream of thermosensors, a plethora of light signaling components have been shown to regulate thermomorphogenesis. Among these, the PHYTOCHROME INTERCTING FACTOR 4 (PIF4) acts as the main hub in regulating thermomorphogenesis (Koini et al., 2009). Other PIFs, including PIF7 also play a role in this process (Fiorucci et al., 2020). Both PIF4 and PIF7 are transcriptionally and post-translationally upregulated by elevated temperature and the stabilized PIF4/7 controls downstream target genes especially auxin biosynthesis and signaling genes to promote hypocotyl elongation (Fiorucci et al., 2020; Franklin et al., 2011; Oh et al., 2012; Stavang et al., 2009). Consistently, all factors affecting PIF4 transcriptionally and/or posttranslationally are involved in thermomorphogenesis. These include an E3 ligase BLADE ON PETIOLE (BOP) that degrades PIF4 post-translationally (Zhang et al., 2017); the DEETIOLATED 1 (DET1)-CONSTITUTIVE PHOTOMORPHOGENIC 1 (COP1)-ELONGATED HYPOCOTYL 5 (HY5) module that controls PIF4 both transcriptionally and post-translationally (Delker et al., 2014; Gangappa and Kumar, 2017), and HEMERA (HMR) that directly interact with PIF4 to stabilize PIF4 and also controls PIF4 target genes to regulate thermomorphogenesis (Qiu et al., 2019).

Apart from light signaling genes, circadian clock and flowering time genes are also involved in regulating thermomorphogenesis. For example, EARLY FLOWERING 3 (ELF3) physically interacts with PIF4 and controls the activity of PIF4 to regulate thermomorphogenesis independent of the circadian clock (Box et al., 2015; Nieto et al., 2015). TIMING OF CAB EXPRESSION 1 (TOC1)-PIF4 interaction results in inhibition of PIF4 function to mediate circadian gating of thermomorphogenic growth at elevated temperature (Zhu et al., 2016). A GIGANTEA (GI)- REPRESSOR OF *ga1-3* (RGA)-PIF4 signaling module enables daylength-dependent modulation of thermomorphogenesis (Park et al., 2020). Furthermore, transcription factors B-box 18 (BBX18) and BBX23 promote thermomorphogenesis via inhibiting ELF3, resulting in activation of PIF4 function (Ding et al., 2018). The RNA-binding protein FCA directly interacts with PIF4 and inhibits the promoter occupancy of PIF4 to attenuate auxin-induced stem elongation at elevated temperature (Lee et al., 2014). Finally, a group of TEOSINTE BRANCHED 1/CYCLOIDEA/PCF (TCP) transcription factors physically interacts with PIF4 and Cry1 to regulate thermomorphogenesis (Han et al., 2019; Zhou et al., 2019). Overall, a number of regulatory proteins are impinging upon PIF4, highlighting the importance of PIF4 in regulating thermomorphogenesis.

The *SUPPRESSOR OF PHYA-105* (*SPA*) family of genes were originally discovered as suppressors of phyA signaling pathways (Hoecker, 2017; Hoecker et al., 1999; Laubinger et al., 2004) and has been shown to function as part of a complex with COP1 as E3 Ubiquitin ligases and play both negative and positive roles in regulating photomorphogenesis (Hoecker, 2017; Pham et al., 2018). Very recently, SPA1 has been shown to function as a ser/thr kinase for the light-induced degradation of PHYTOCHROME INTERACTING FACTOR 1 (PIF1) (Paik et al., 2019). Although COP1-SPA complex has been shown to function in thermomorphogenesis (Delker et al., 2014; Park et al., 2017), the molecular functions of SPA proteins and the significance of the SPA kinase activity in high ambient temperature signaling have not been shown yet. Here, we show that the SPA proteins as well as the SPA1 kinase activity are essential for regulating thermomorphogenesis via controlling the phyB-PIF4 module.

## RESULTS AND DISCUSSION

### *SPA*s are required for thermomorphogenesis

To thoroughly examine whether the *SPA* genes are required for thermomorphogenesis, we tested high ambient temperature mediated hypocotyl elongation phenotype using *spa* higher order mutants and TAP tagged *SPA1* overexpression plants driven by the CaMV 35S promoter (Figure 1A and B). To this end, we grew seedlings at 22°C under continuous white light for two days and then treated either at 22°C or at 28°C for additional four days. While we could not observe statistically significant difference in hypocotyl growth in 22°C for *spa* double and triple mutants, the higher order *spa* mutants exhibited shorter hypocotyl in high ambient temperature (28°C). Interestingly, *spaQ* mutant in which all four *SPAs* were mutated, displayed insensitivity to high ambient temperature treatment, indicating that *SPA*s are necessary for the thermomorphogenesis in a redundant manner. However, the overexpressed *SPA1* (*35S:TAP-SPA1*) line showed similar phenotype with wild-type, indicating that an excessive amount of SPA1 may not necessarily enhance the high ambient temperature mediated hypocotyl elongation. We then examined whether *SPA1* can complement the thermomorphogenic response in *spaQ* mutant. We used previously reported *LUC-SPA1/spaQ* transgenic plants (Paik et al., 2019). As expected, SPA1 can partially rescue the *spaQ* mutant thermo-insensitive phenotype (Figure 1A and B), indicating that SPA1 is required for thermomorphogenesis. To investigate the potential temperature regulation on SPA1, we first examined SPA1 protein level at 22°C and 28°C using *35S:TAP-SPA1* plant (Figure 1C and D). Intriguingly, SPA1 protein level did not change at high ambient temperature indicating that temperature does not post-translationally regulate SPA1 protein stability. We also examined the transcript level of *SPA*s at high ambient temperature (Figure S1), and did not observe any difference in gene expression of *SPAs*, except *SPA1* which showed lower expression at high ambient temperature. Taken together, our results showed that *SPA*s are required for thermomorphogenic hypocotyl growth.

**Figure 1:**
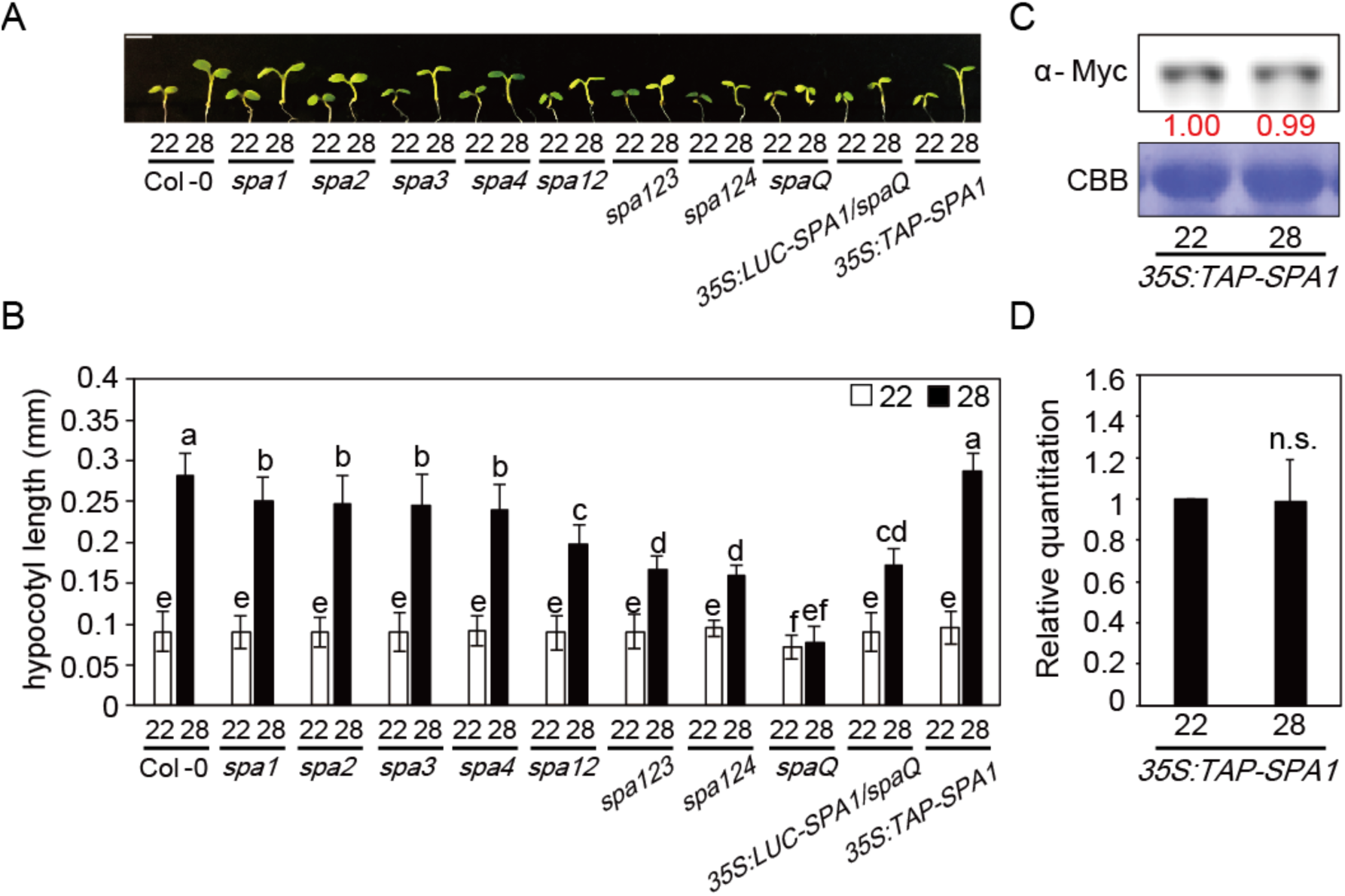
SPAs are required for thermomorphogenesis. (A) Photograph showing seedling phenotypes of *spa* high order mutants in normal and high ambient temperature. Seedlings were grown for two days in continuous white light at 22°C and then either kept at 22°C or transferred to 28°C for additional four days before being photographed. More than 10 seedlings were measured for each experiment and was repeated 3 times for one-way ANOVA analysis. The scale bar represents 2 mm. (B) Bar graph shows the hypocotyl lengths for seedlings grown under conditions described in (A). The letters a-f indicate statistically significant differences between means of hypocotyl lengths (P<0.05) based on one-way ANOVA analyses. Error bars indicate s.d. (n=3). (C) Western blot shows the level of TAP-SPA1 from *35S:TAP-SPA1* whole seedlings grown for 5 days in 22°C and either kept at 22°C or transferred to 28°C for 4 hours. TAP-SPA1 was detected using Myc antibody. Red number indicates the quantitation value from anti-Myc band divided by the Coomassie staining band intensity. (D) Bar graph shows the relative amount of TAP-SPA1 (n=3).

### *SPA*s control global gene expression at high ambient temperature

High ambient temperature induces global changes in gene expression to affect thermomorphogenesis (Jung et al., 2016). To test if SPAs can regulate gene expression in response to high ambient temperature, RNA-seq was conducted using wild-type Col-0 and *spaQ* mutant at both 22°C and 28°C. The results show that 4470 genes were differentially regulated in wild-type whereas only 1379 genes were regulated in *spaQ* mutant in response to high ambient temperature (Figure 2A). Moreover, 735 genes were shared between the wild type and *spaQ*, which is >53% of genes regulated in *spaQ*. A closer examination showed that >54% of up-regulated genes and 38.5% of down-regulated genes in *spaQ* were shared with wild-type (Figure 2A), indicating that *SPA*s are crucial for transcriptional regulation in thermomorphogenesis. We analyzed 2088 genes which are only up-regulated in wild-type but were not affected in *spaQ* mutant, and also 1726 genes which are only down-regulated in wild-type but were not altered in *spaQ* mutant using Gene Ontology (GO) analysis (Figure 2B and C). Many of these genes were responsible for abiotic responses such as water and stress responses in both wild-type up-regulated and down-regulated genes. Notably, many of the detoxification and secondary metabolite biosynthesis responsive genes were involved in wild-type thermomorphogenesis, but not in *spaQ* mutant. Also, many genes which are involved in metabolism or development such as photosynthesis, cell maturation or root development were down-regulated in wild-type but not in *spaQ* mutant. Furthermore, 735 DEGs were shared in both wild-type and *spaQ* which displayed four distinct patterns as shown in the heatmap analysis (Figure 2D). Although the expression levels are different, two groups (I and IV) displayed similar pattern between the wild-type and *spaQ* mutant, whereas the other two (II and III) displayed opposite pattern of expression. We further conducted qPCR to confirm the RNA-seq data using 3 different genes including genes from group II and III (Figure 2E). *SAUR20*, which is one of the 2088 genes in Fig. 2B, showed similar expression pattern in both qPCR and RNAseq analyses, which is upregulated in wild-type but did not change in *spaQ* mutant at high ambient temperature. *At4g33720* from group II and *At1g09380* from group III also displayed similar pattern which show opposite regulation between the wild-type and *spaQ* mutant. Finally, we examined that the transcript level of several thermo-responsive marker genes such as *IAA 29, PRE 5, SAUR 15*, and *CYP79B2*. The expression of these genes was not up- or down-regulated in *spaQ* mutant, whereas obvious regulation was observed in wild type (Figure S2). Overall, these data suggest that *SPA*s play a crucial role in thermomorphogenesis through transcriptional regulation.

**Figure 2:**
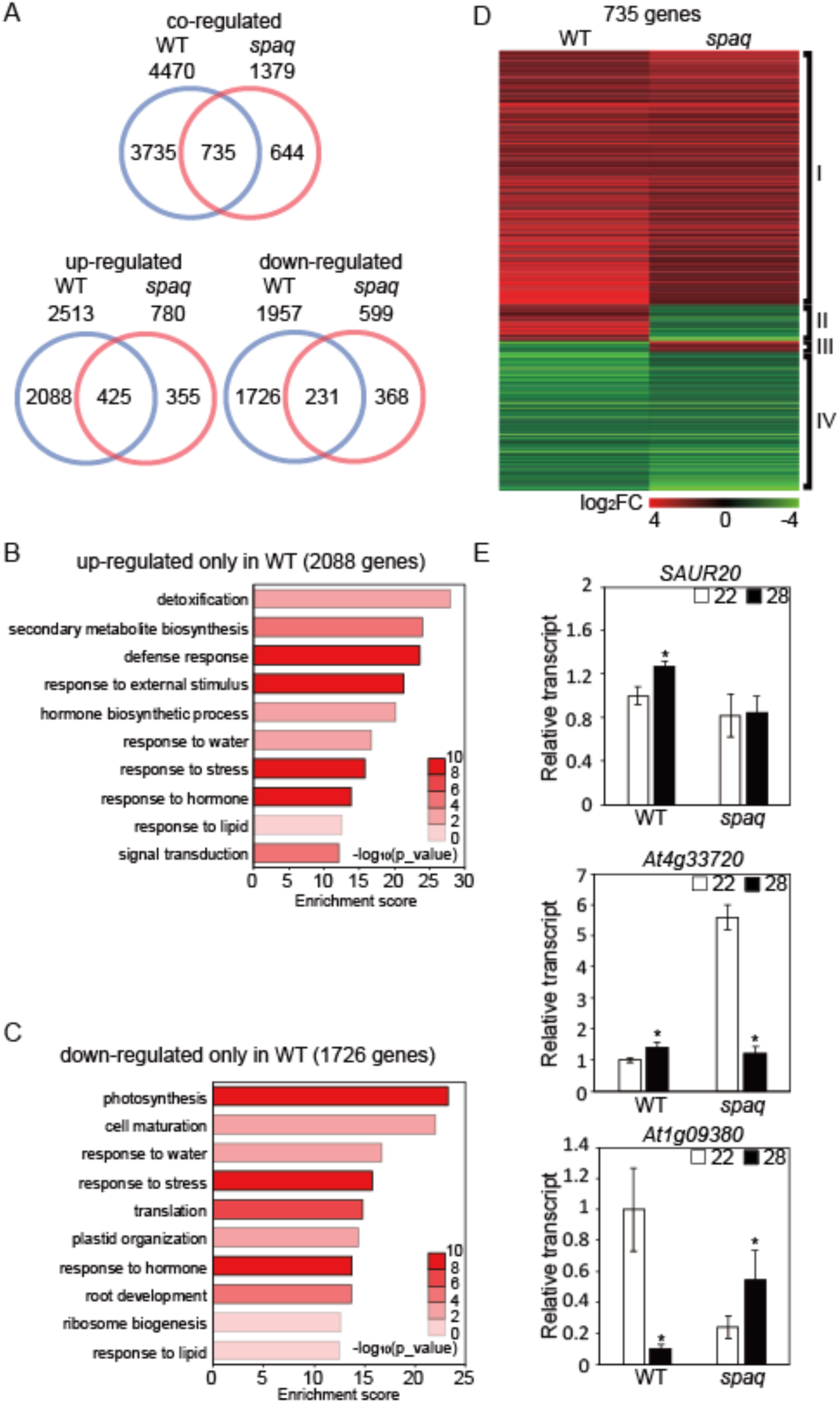
SPAs regulate global gene expression at high ambient temperature. (A) Venn diagrams show co-regulated, up-regulated, and down-regulated genes in wild-type (WT) vs *spaQ* mutant at high ambient temperature response. Six-day-old white light-grown seedlings were transferred to 22°C or 28°C for additional 24 hours and total RNA was extracted from three biological replicates for RNA-seq analyses. (B) Gene Ontology (GO) analysis of up-regulated genes only in WT. (C) Gene Ontology (GO) analysis of down-regulated genes only in WT. (D) Hierarchical clustering displaying 735 differentially expressed genes from co-regulated genes shared between wild type and *spaQ* as shown in (A). Co-regulated genes were identified with FDR<0.05. Each group represents up-regulated in both WT and *spaQ* (I), up-regulated in WT but down-regulated in *spaQ* (II), down-regulated in WT but up-regulated in *spaQ* (III), and down-regulated in both WT and *spaQ* (IV) respectively. (E) RT-qPCR analysis using *SAUR20*, already identified as high ambient temperature marker gene, *At4g33720* from group II from (D), *At1g09380* from group III from (D). RT-qPCR samples were from Col-0 whole seedlings grown for 5 days at 22°C and then either kept at 22°C or transferred to 28°C for 24 hours. Three biological repeats were performed. Relative gene expression levels were normalized using expression levels of *ACT7*. Asterisks indicate statistically significant difference using Student’s t-test; *p < 0.05, **p < 0.01.

### SPAs are essential for PIF4 stabilization at high ambient temperature

PIF4 is considered as a main hub in high ambient temperature response pathways (Paik et al., 2017; Pham et al., 2018). The expression and stability of PIF4 are regulated under high ambient temperature (Foreman et al., 2011; Oh et al., 2012; Stavang et al., 2009). Stabilized PIF4 triggers transcriptional regulation in auxin responsive and BR biosynthetic genes to promote thermomorphogenesis (Franklin et al., 2011; Jung et al., 2016; Koini et al., 2009; Legris et al., 2019; Martínez et al., 2018; Oh et al., 2012; Qiu et al., 2019). Since *spaQ* mutant did not respond to high ambient temperature-induced hypocotyl elongation, we examined whether SPAs are regulating the thermomophogenesis through PIF4 at high ambient temperature. To test genetic relationship between PIF4 and SPAs, we generated *35S:PIF4-Myc* overexpression line in *spaQ* mutant background and measured the phenotype at high ambient temperature (Figure 3A and B). Results show that the hypocotyl lengths of *35S:PIF4-Myc/spaQ* and *spaQ* were similar at high ambient temperature, suggesting that *spaQ* is epistatic to *PIF4* overexpression phenotype. To examine if SPA proteins regulate PIF4 protein stability, we performed immunoblot analyses from *35S:PIF4-Myc* and *35S:PIF4-Myc*/*spaQ* grown at 22°C and 28°C. Strikingly, PIF4-Myc level was not detectable in *35S:PIF4-Myc*/*spaQ* compared to that in *35S:PIF4-Myc* in wild-type (Figure 3C and D), indicating that SPAs are essential for stabilization of PIF4 at both normal and elevated temperature. We then conducted yeast two hybrid assay using PIF4 and SPA1 to examine whether these proteins interact directly. The result show that PIF4 interacts with SPA1 in yeast (Figure S3A). In addition, we also confirmed the interaction between SPA1 and PIF4 through *in vitro* pull-down assay using bacterially expressed MBP tagged SPA1 and GST tagged PIF4 (Figure S3B). These data suggest that SPA1 might regulate PIF4 via physical interaction. Taken together, these data suggest that SPAs might promote thermomorphogenesis under high ambient temperature through stabilization of PIF4.

**Figure 3:**
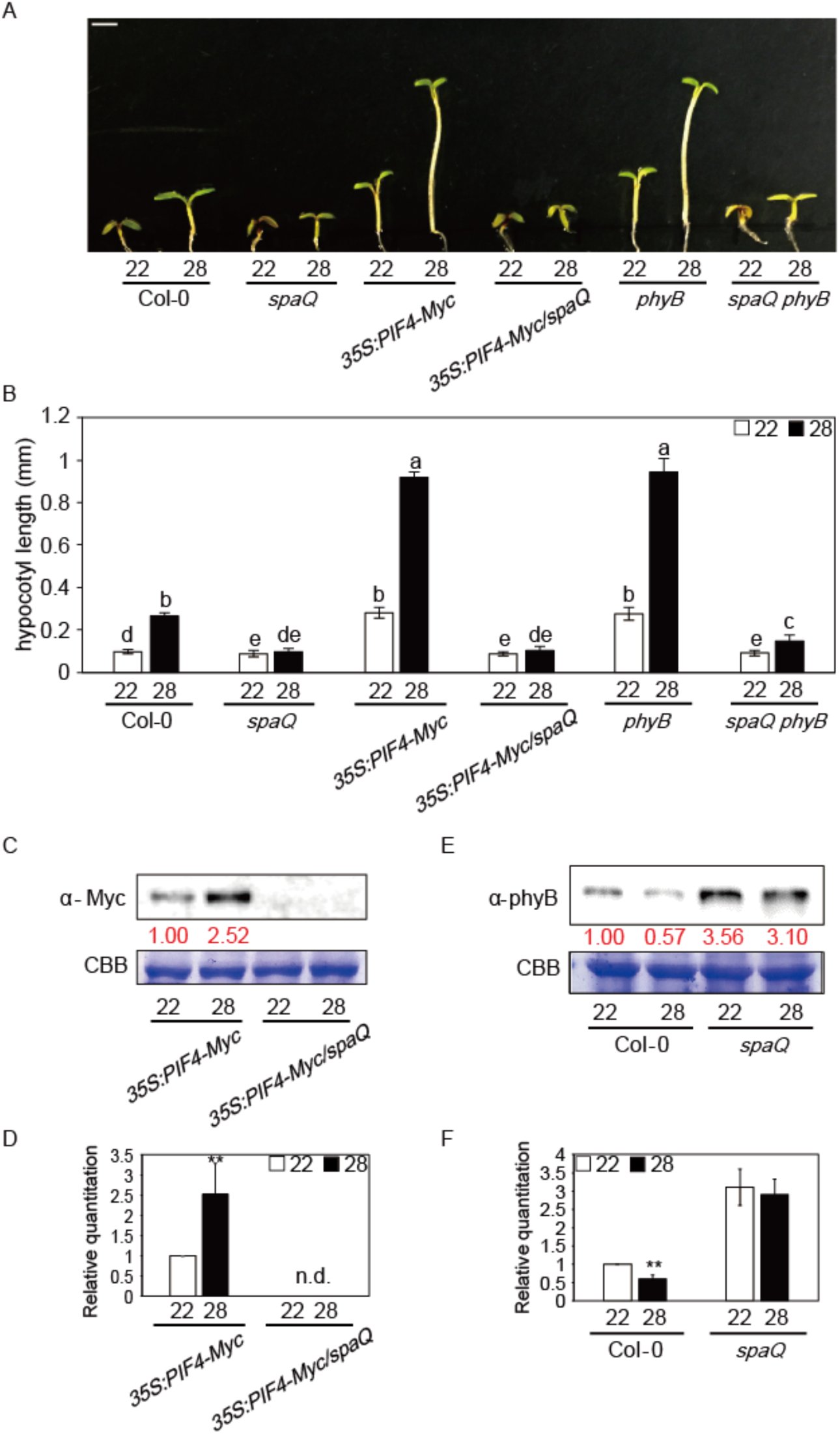
SPAs are necessary for controlling the phyB-PIF4 module at high ambient temperature. (A) Photograph shows the seedling phenotypes of Col-0, *spaQ, 35S:PIF4-Myc, 35S:PIF4-Myc*/*spaQ, phyB*, and *spaQ phyB* plants at 22°C or 28°C. Seedlings were grown for two days in continuous white light at 22°C and then either kept at 22°C or transferred to 28°C for additional four days before being photographed. More than 10 seedlings were measured for each experiment and was repeated 3 times for one-way ANOVA analysis. The scale bar represents 2 mm. (B) Bar graph shows the hypocotyl lengths for seedlings grown under conditions described in (A). The letters a-e indicate statistically significant differences between means of hypocotyl lengths (P<0.05) based on one-way ANOVA analyses. Error bars indicate s.d. (n=3). (C) Western blot shows PIF4-Myc protein level from either *35S:PIF4-Myc* or *35S:PIF4-Myc*/*spaQ* whole seedling grown for 5 days in 22°C and transferred to 22°C or 28°C for 4 hours. PIF4-Myc was detected using Myc antibody. Red number indicates the quantitation value from anti-Myc detection divided by Coomassie blue staining intensity. (D) Bar graph shows the relative amount of PIF4-myc (n=3). n.d. = not detectable. (E) Western blot shows the phyB protein level from either Col-0 or *spaQ* mutant at 22°C or 28°C. (F) Bar graph shows the relative amount of phyB (n=3). Asterisks indicate statistically significant difference using Student’s t-test; **p < 0.01.

### SPAs are essential for phyB degradation at high ambient temperature

Phytochrome B (phyB) has been shown to function as a temperature sensor not only under diurnal conditions but also under continuous light (Jung et al., 2016; Legris et al., 2016; Qiu et al., 2019). Phytochromes are well-known red/far-red-light receptors (Legris et al., 2019), and SPAs are one of the key components in light signaling pathways that bind to phyB (Lu et al., 2015; Paik et al., 2019; Sheerin et al., 2015). Since phyB and SPA1 have been shown to interact directly with each other, we hypothesized phyB and SPAs might also be tightly linked in temperature signaling cascade. Therefore, we measured phyB level in WT and *spaQ* mutant using phyB antibody. Interestingly, we first observed that phyB level was decreased in wild type at high ambient temperature. However, unlike wild type, phyB level was elevated at 22°C and remained similar in *spaQ* mutant at high ambient temperature (Figure 3E and F). These data suggest that high ambient temperature induces thermomorphic response by decreasing the phyB protein level possibly due to receptor desensitization, and the SPAs are necessary for this reduction in phyB level. Furthermore, SPAs are required for phyB degradation at both normal and high ambient temperature. We also tested *spaQ phyB* mutant phenotype to examine whether an absence/degradation of phyB has a role in thermomorphogenesis. Interestingly, *spaQ phyB* mutant displayed partially but weakly recovered phenotype compared to *spaQ* at high ambient temperature (Figure 3A and B). These data indicate that the SPA-mediated regulation of phyB level at high ambient temperature is essential for thermomorphogenic hypocotyl elongation.

### Phosphorylation of PIF4 by SPA1 is necessary for thermomorphogenesis

Recently, we reported that SPAs act as ser/thr kinases which phosphorylate PIF1 and control PIF1 stability in response to light (Paik et al., 2019). Since PIF4 also interacts with SPA1, we hypothesized that SPA1 might phosphorylate PIF4. Therefore, we performed an *in vitro* kinase assay to examine whether SPA1 can directly phosphorylate PIF4 (Figure 4A). A strep-tagged full-length SPA1 protein purified from *Pichia pastoris* was used for the kinase assay along with bacterially expressed GST-PIF4. As expected, PIF4 was phosphorylated by SPA1 *in vitro*, indicating that PIF4 might be a *bona fide* substrate of SPA1.

**Figure 4:**
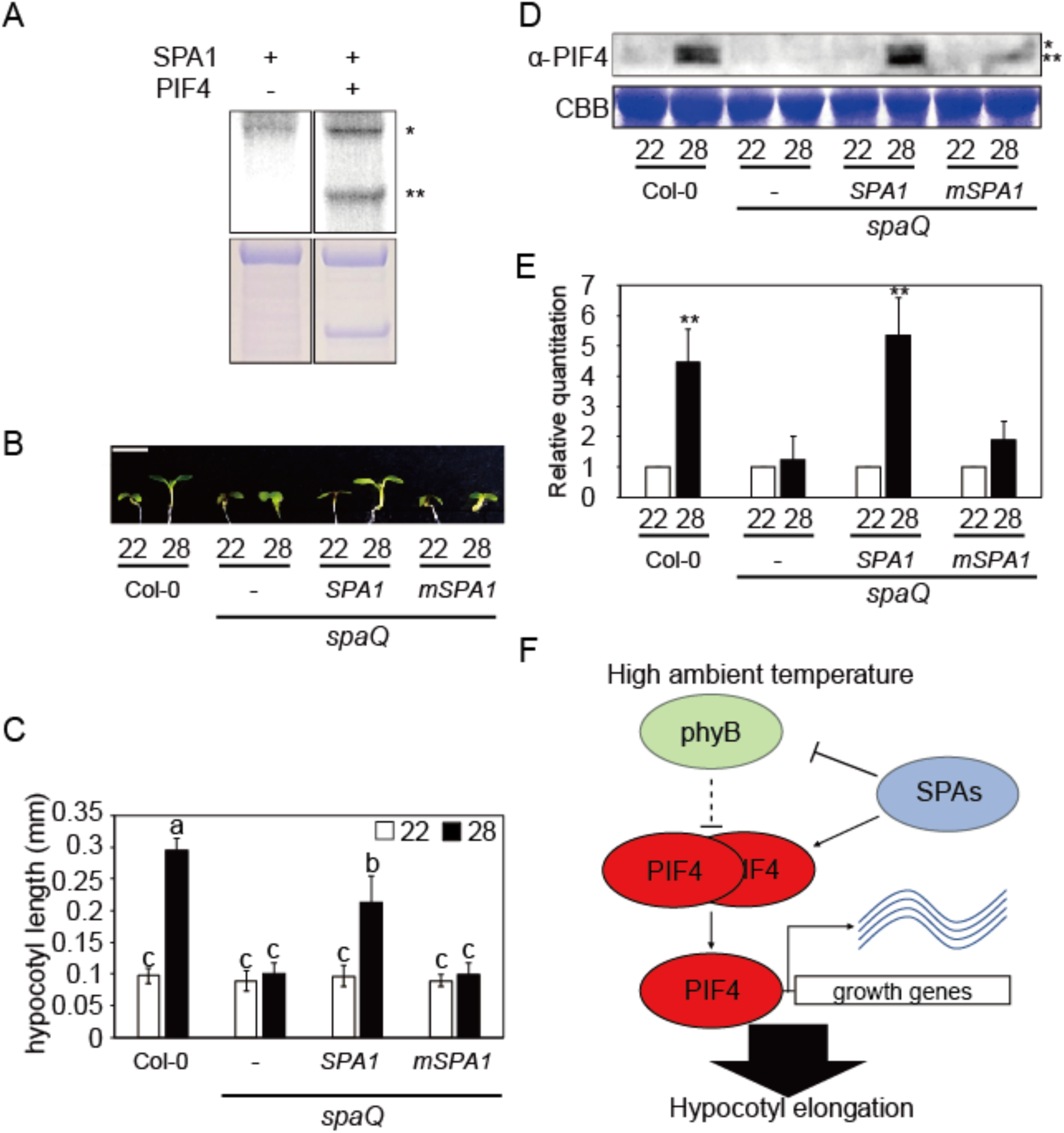
Phosphorylation of PIF4 by SPA1 is essential for thermomorphogenesis. (A) (Top) Autoradiograph shows that SPA1 directly phosphorylates PIF4 *in vitro. In vitro* kinase assay performed using purified SPA1 from *Pichia pastoris* and GST PIF4 from *E*.*coli*. Upper single asterisk (*) shows SPA1 band and lower double asterisks (**) shows PIF4 band. (Bottom) Coomassie-stained gel shows the amount of proteins used in the assay. (B) Photograph shows the seedling phenotypes of Col-0, *spaQ, 35S:LUC-SPA1*/*spaQ*, and *35S: LUC-mSPA1*/*spaQ* at 22°C or 28°C. Three *spaQ* background plants are indicated with - for *spaQ, SPA1* for *35S:LUC-SPA1*/*spaQ*, and *mSPA1* for *35S:LUC-mSPA1*/*spaQ*, respectively. Seedlings were grown for two days in continuous white light at 22°C and then either kept at 22°C or transferred to 28°C for additional four days before being photographed. More than 10 seedlings were measured for each experiment and was repeated 3 times for one-way ANOVA analysis. The scale bar represents 2 mm. The letters a-c indicate statistically significant differences between means of hypocotyl lengths (P<0.05) based on one-way ANOVA analyses. Error bars indicate s.d. (n=3). (C) Bar graph shows the hypocotyl lengths for seedlings grown under conditions described in (B). (D) Western blot shows native PIF4 protein level from Col-0, *spaQ, 35S:LUC-SPA1*/*spaQ*, and *35S: LUC-mSPA1*/*spaQ* whole seedling grown for 5 days at 22°C and then either kept at 22°C or transferred to 28°C for 4 hours. PIF4 was detected using native PIF4 antibody. Upper single asterisk (*) shows phosphorylated form and lower double asterisks (**) shows unphosphorylated PIF4. (E) Bar graph shows the relative amount of PIF4 (n=3). Asterisks indicate statistically significant difference using Student’s t-test; **p < 0.01. (F) Simplified model shows the role of SPA proteins in regulating thermomorphogenesis under elevated temperature. After high ambient temperature exposure, SPAs promote the degradation of phyB, but stabilize PIF4 either by direct phosphorylation and/or other unknown mechanisms. Stabilized PIF4 then promotes hypocotyl elongation through direct promoter binding to hypocotyl elongation responsive genes.

A conserved amino acid mutation (R517E) in the kinase domain of SPA1 greatly reduces its kinase activity *in vitro* (Paik et al., 2019). Thus, we measured the phenotype of *35S:LUC-mSPA1*/*spaQ* at high ambient temperature to examine whether the kinase domain of SPA1 is necessary for thermomorphic hypocotyl elongation (Figure 4B, C). In contrast to wild type *35S:LUC-SPA1*/*spaQ, 35S:LUC-mSPA1*/*spaQ* did not rescue the thermo-insensitive phenotype of *spaQ* at high ambient temperature indicating that the kinase activity of SPA1 is necessary for thermomorphogenesis. Moreover, an immunoblot using anti-PIF4 antibody showed that both the phosphorylated and unphosphorylated forms of native PIF4 are unstable in the *spaQ* background compared to wild type (Figure 4D, E). Strikingly, the PIF4 level was not rescued in the *35S:LUC-mSPA1*/*spaQ*, while the expression of *35S:LUC-SPA1*/*spaQ* largely rescued both the phosphorylated and unphosphorylated forms of PIF4 consistent with the phenotype of the *35S:LUC-SPA1*/*spaQ* seedlings. Taken together, these data suggest that SPAs might regulate thermomorphogenesis through direct phosphorylation of PIF4. Previously, the BR signaling kinase BRASSINOSTEROID-INSENSITIVE 2 (BIN2) has been shown to directly phosphorylate PIF4, and the BIN2-mediated phosphorylation destabilizes PIF4 (Bernardo-García et al., 2014). In contrast, SPA proteins stabilize PIF4. Thus, multiple antagonistic kinases might fine tune PIF4 level to regulate seedling development under ambient and elevated temperatures.

In summary, our data uncovered a new role of SPA proteins that act as positive regulators of thermomorphogenesis by controlling the phyB-PIF4 module (Figure 4D). While SPAs destabilize the thermosensor phyB, they are crucial for stabilizing PIF4, a main hub of thermomorphogenesis. In contrast to SPAs, ELF3, TOC1, FCA, TCP and others interact with PIF4 and control the activity of PIF4 to regulate thermomorphogenesis (Lee et al., 2014; Nieto et al., 2015; Paik et al., 2017; Zhou et al., 2019; Zhu et al., 2016). Thus, SPAs in concert with other factors fine tune thermomorphogenesis in response to high ambient temperature.

## MATERIALS AND METHODS

### Plant materials, growth conditions and phenotypic analyses

Seeds from Col-0 ecotype of *Arabidopsis thaliana* were used in this study. Seeds were surface-sterilized and plated on no sucrose contained Murashige and Skoog (MS) medium. After 3 days at 4 for stratification, seedlings were placed in 22°C for 2 days and transferred to 22°C or 28°C for 4 additional days. Hypocotyl length of more than 15 seedlings were measured using ImageJ software and statistically analyzed using one-way ANOVA analysis.

### Protein extraction and Western blot analyses

Total protein of 50 seedlings for each sample was extracted using 50 uL urea extraction buffer [8 M urea, 0.35 M Tris-Cl pH 7.5, 1 mM phenylmethylsulfonyl fluoride (PMSF) and 1× protease inhibitor cocktail]. Subsequently, 6X SDS buffer were added followed by 5 min of denaturation at 95°C. Then, samples were centrifuged at 16,000 g for 15 min. Supernatant from the samples was loaded into 8% SDS-PAGE gels for separation. PVDF membrane (Millipore) were used for transfer, and Western blots were detected using anti-Myc (cell signaling) or anti-phyB antibody. Coomassie blue staining was also used for the loading control.

### RNA extraction, cDNA synthesis, and qRT-PCR

RNA-Seq was performed using 6-day-old white light-grown seedlings with three independent biological replicates (n = 3). Seeds were kept in 22°C under continuous white light for 6 days. After 6 days, seedlings were either kept at 22°C or transferred to 28°C for additional 24 hours under continuous white light. Total RNA was extracted from these seedlings using Plant RNA purification kit (Sigma-Aldrich) according to the manufacturer’s protocols. For cDNA synthesis, 2 μg of total RNA was used for reverse transcription with M-MLV Reverse Transcriptase (Thermo Fisher Scientific). SYBR Green PCR master mix (Thermo Fisher Scientific) and gene-specific oligonucleotides (Key Resources Table) were used to conduct qPCR analyses. Finally, relative transcription level was calculated using 2^ΔCt^ with *ACT2* normalization.

### RNA-seq analyses

For the RNA-seq analysis, 3’Tag-Seq method was used in this study (Lohman et al., 2016). Raw read quality was accessed using FastQC (www.bioinformatics.babraham.ac.uk/projects/fastqc/). Then raw reads were aligned to the Arabidopsis genome using Bowtie2 (Langmead and Salzberg, 2012) and TopHat (Trapnell et al., 2012). The annotation of the Arabidopsis genome was obtained from TAIR10 (www.arabidopsis.org/). Read count data were performed by HTseq (Anders et al., 2015) (htseq.readthedocs.io/en/master/index.html). Differentially expressed genes in *spaQ*/WT were identified using the EdgeR (Robinson et al., 2010). The differential gene expression was defined using cutoff of ≥2-fold with adjusted P value (FDR) ≤0.05. Venn diagrams were generated using the website (http://bioinformatics.psb.ugent.be/webtools/Venn/). Heatmap was generated using Morpheus (https://software.broadinstitute.org/morpheus/). We used the options as Hierarchical clustering with one minus cosine similarity metric combined with average linkage method for the heatmap. Also, GO enrichment analyses were performed using TAIR (https://www.arabidopsis.org/tools/go_term_enrichment.jsp). GO bar graphs were generated based on the significant enriched terms with the lowest P value and FDR (≤0.05) for GO terms.

### Yeast two-hybrid analyses

Cloning of *SPA1* in pJG4-5 have been described previously (Xu et al., 2014). *PIF4* was cloned into pEG202 vector using EcoRI and XhoI restriction enzyme sites. Different combinations of AD-LexA and BD-fusion plasmids were introduced into yeast strain EGY48-0 and selected on -His, -Ura, -Trp minimal synthetic medium. β-galactosidase assay was performed according to the manufacturer’s protocol (Matchmaker Two-Hybrid System; Takara, https://www.takarabio.com).

### Pull-down assays

For *in vitro* pull-down assays, MBP-SPA1 (Xu et al., 2014) and GST-PIF4 were used in this study. GST-PIF4 was expressed from pGEX4T-1 vector. Bacterial extract expressing GST-PIF4 was incubated with glutathione resin in the binding buffer (50 mM Tris-Cl pH7.5, 200 mM NaCl, 0.1% NP-40) for 2 hours. Resin was washed more than 5 times with wash buffer (1x PBS, 0.1% Tween 20). Samples were boiled and analyzed using Western blot. Anti-MBP (E8032S, New England Biolabs) and Anti-GST-HRP conjugate (RPN1236; GE Healthcare Bio-Sciences) were used to detect MBP-SPA1 and GST or GST-PIF4, respectively.

### In-vitro kinase assay

For SPA1 kinase assay, approximately 500 ng of SPA1 and 1 μg of GST-PIF4 were used using the kinase buffer (50 mM Tris, pH 7.5, 4 mM β-mercaptoethanol, 1 mM EDTA, 10 mM MgCl2). ^32^P radio-labeled γ-ATP (Perkin Elmer Cat# BLU502A) was added to the reaction and incubated at 28°C for 1 h. 6x SDS sample buffer was used to stop the reaction and boiled proteins were loaded to 7.8% SDS-PAGE gel. After separation, gels were dried and exposed to a phosphor screen and then scanned using Typhoon FLA 9500 (GE healthcare).

## ACKNOWLEDGEMENT

We thank Dr. Sibum Sung and the members of the Huq laboratory for critical reading of the manuscript, Drs. Hong-Quan Yang for sharing *spaQ phyB* seeds and Peter Quail for sharing anti-phyB antibody, Dr. Taeyoung Lee for comments analyzing RNA-seq. The authors acknowledge the Texas Advanced Computing Center (TACC) at The University of Texas at Austin for providing High Performance Computing, visualization, and database resources that have contributed to the research results reported in this paper.

## AUTHOR CONTRIBUTION

S.L, I.P. and E.H. conceived the study and designed the experiments. S.L and I.P carried out the experiments. S.L, I.P. and E.H. analyzed the data. S.L. and E.H. wrote the paper and I.P. commented on the paper.

## DATA AVAILABILITY

Raw data and processed data for RNA-Seq in Col-0 and *spaQ* can be accessed from the Gene Expression Omnibus database under accession number GSE142354.

## Figure legends

**Figure S1:**
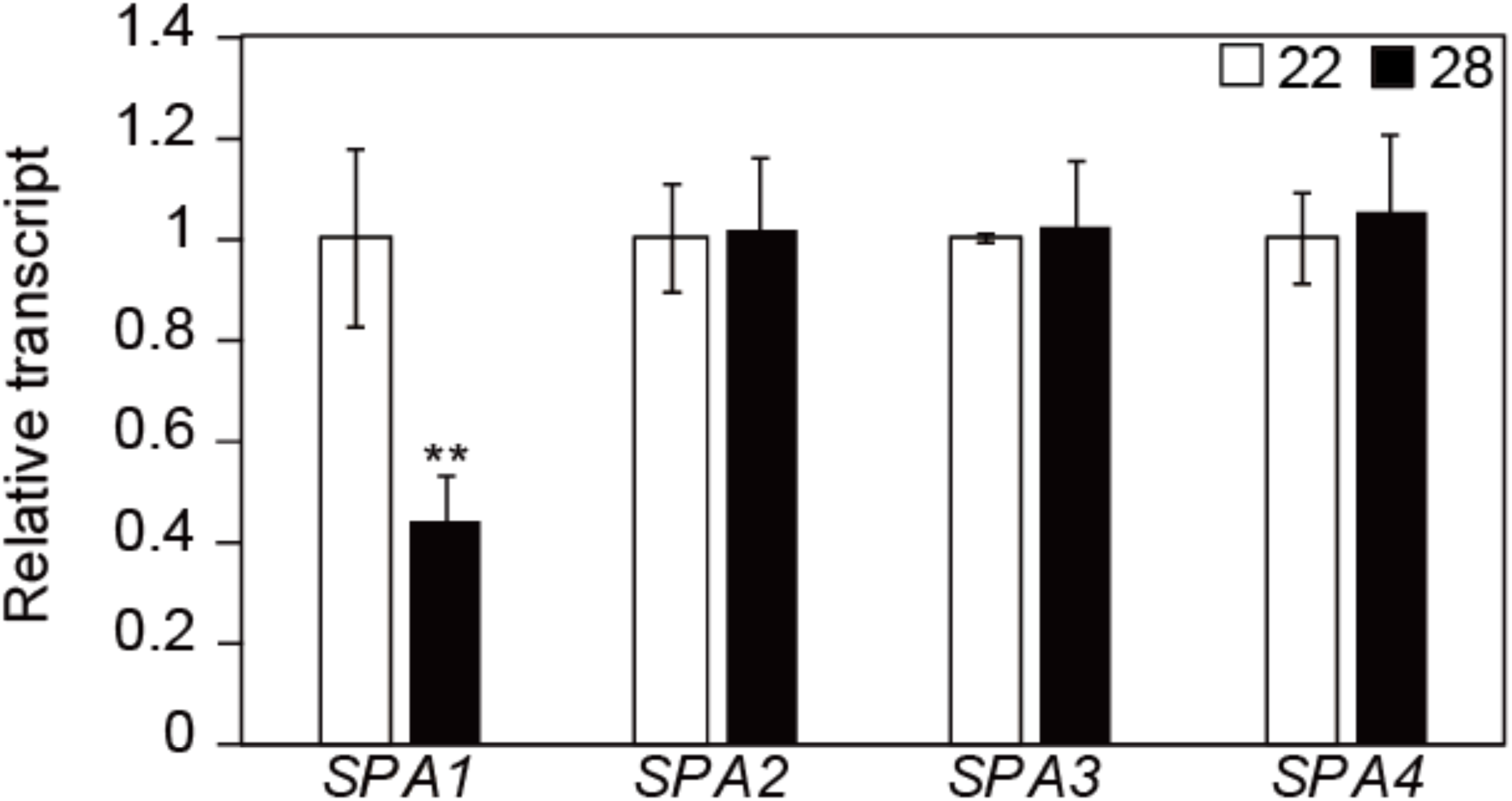
Relative transcript level of *SPA*s. qPCR was performed to detect 4 *SPA*s transcriptional level. Samples were from Col-0 whole seedling grown for 5 days in 22°C and transferred to 22°C or 28°C for 4 hours. Three biological replicates were used in this study. Relative gene expression levels were normalized using expression levels of *ACT7*. Asterisks indicate statistical difference using Student’s t-test; **p < 0.01.

**Figure S2:**
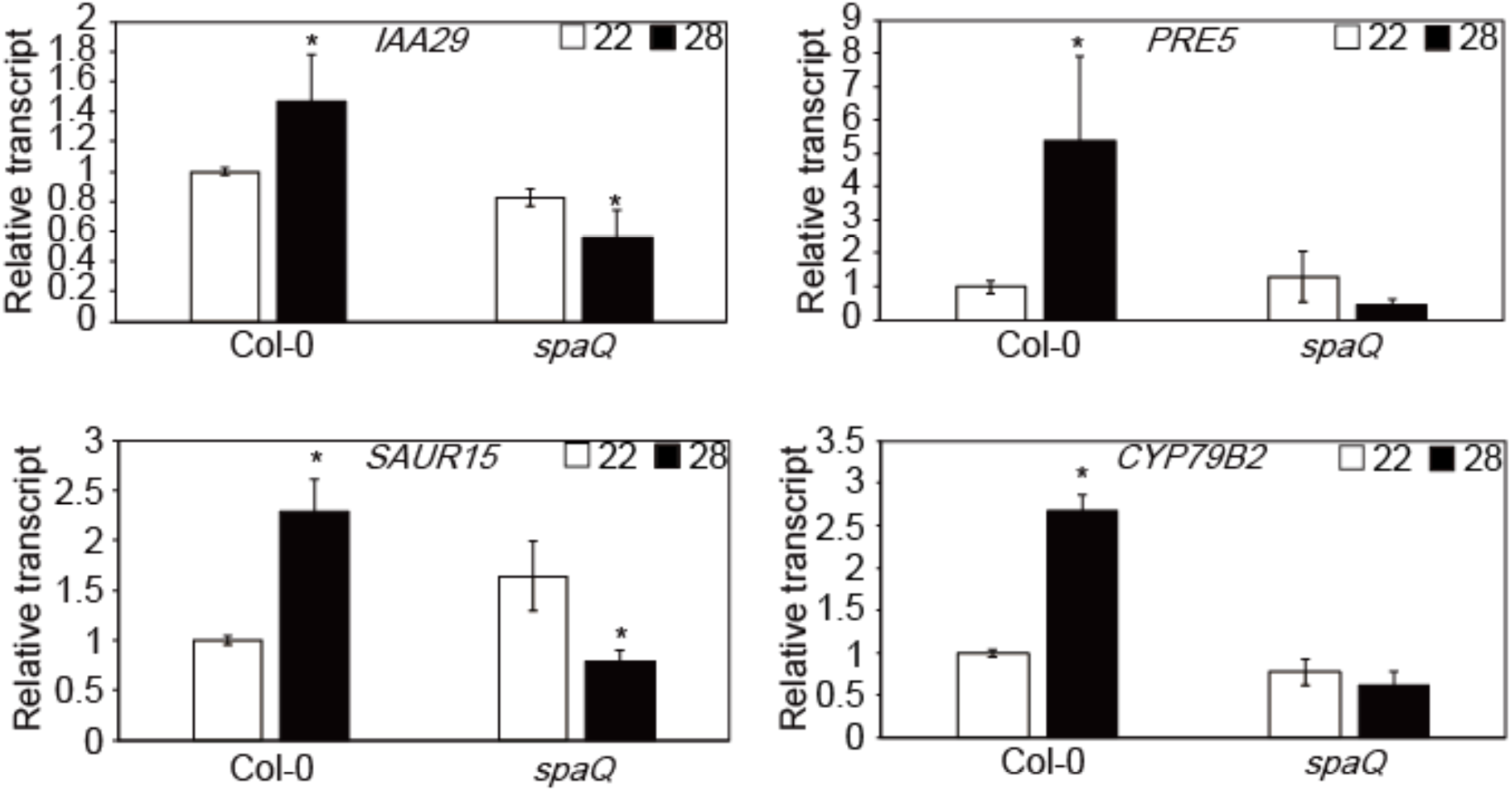
Relative transcript level of thermo-responsive genes. qPCR was performed to detect transcriptional level of *IAA 29, PRE 5, SAUR 15*, and *CYP79B2*. Samples were from Col-0 or *spaQ* mutant whole seedling grown for 5 days in 22°C and transferred to 22°C or 28°C for 4 hours. Three biological replicates were used in this study. Relative gene expression levels were normalized using expression levels of *ACT7*. Asterisks indicate statistical difference using Student’s t-test; *p < 0.05.

**Figure S3:**
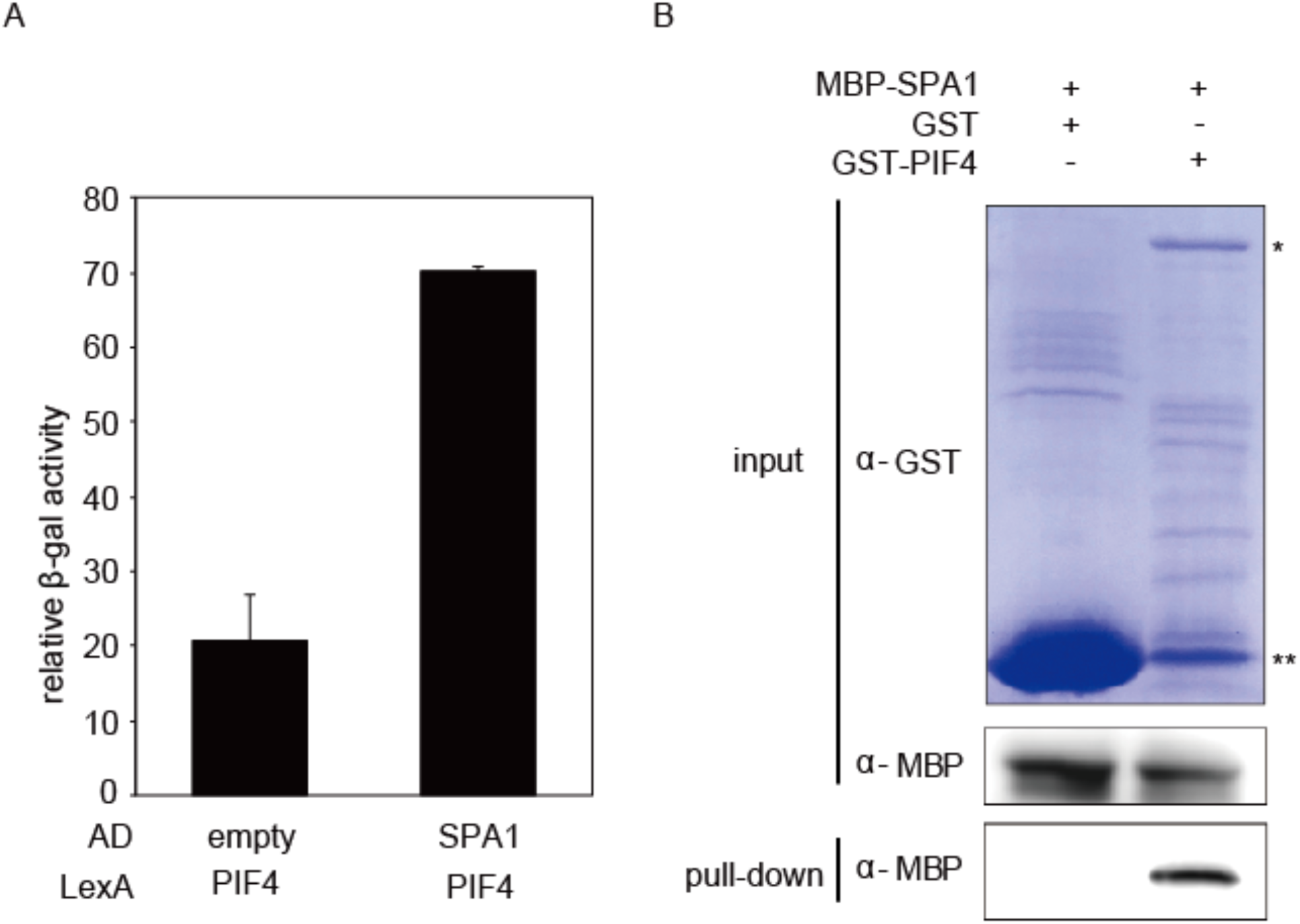
Interaction between SPA1 and PIF4. (A) yeast two hybrid of SPA1 and PIF4. LexA-PIF4 was co-transformed with empty AD or AD-SPA1. The error bars represent standard deviation. Three biological replicates were used in this study. (B) *in vitro* pull-down assay using MBP-SPA1 and GST-PIF4. GST only was used for negative control. Single asterisk mark (*) shows GST-PIF4 band and double asterisk mark (**) shows GST only protein.

**Figure S4:**
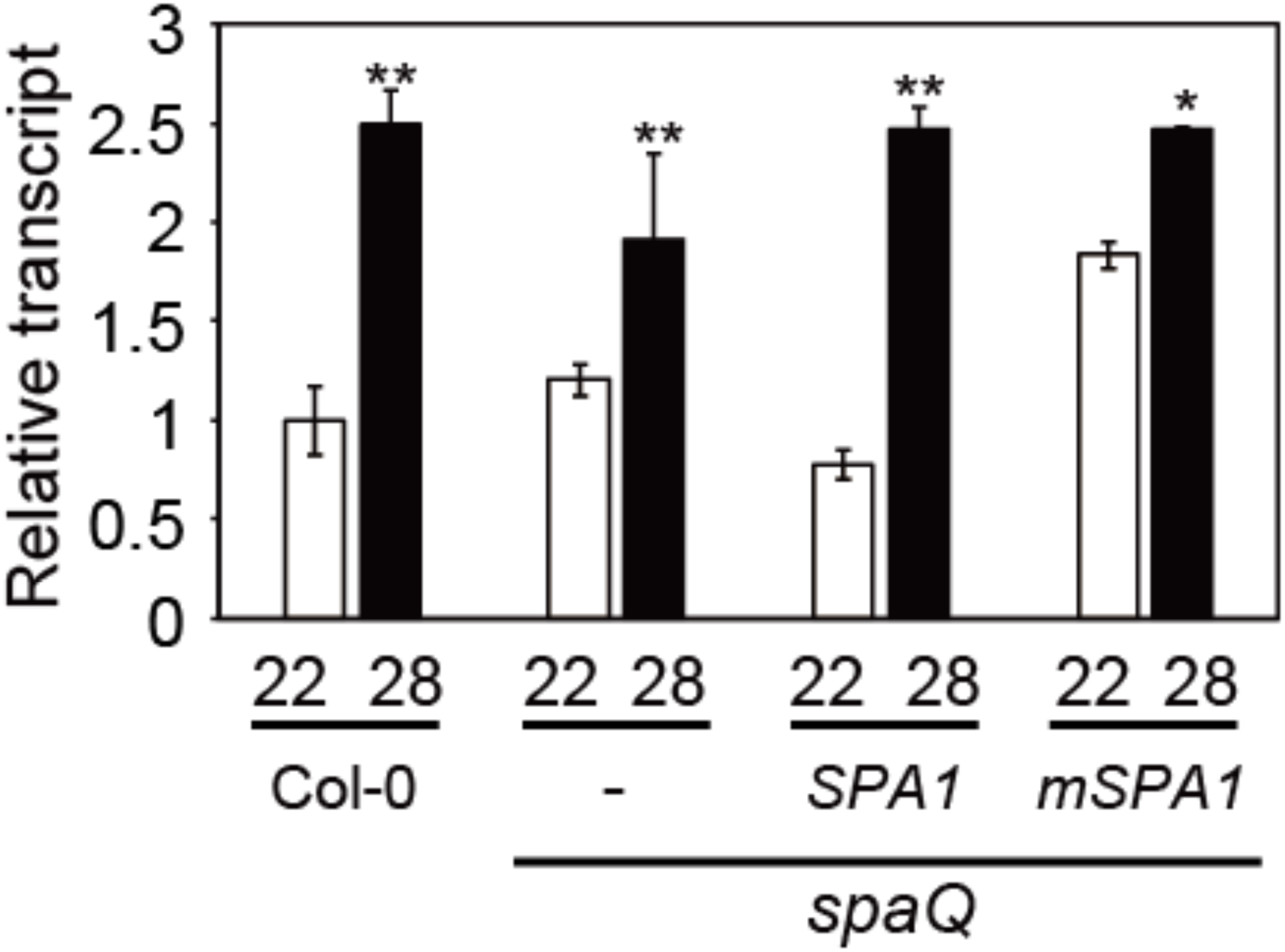
Relative expression level of *PIF4*. Transcription level of *PIF4* using qPCR analysis. qPCR samples were from Col-0, *spaQ, 35S:SPA1-LUC/spaQ*, and *35S:mSPA1-LUC/spaQ* whole seedling grown for 5 days in 22°C and transferred to 22°C or 28°C for additional 24 hours. Three biological repeats were performed. Relative gene expression levels were normalized using expression levels of *ACT7*. Asterisks indicate statistical difference using Student’s t-test; *p < 0.05, **p < 0.01.

